# Motor Output Matters: Evidence of a Continuous Relationship Between Stop/No-go P300 Amplitude and Response Force on Failed Inhibitions at the Trial-Level

**DOI:** 10.1101/706978

**Authors:** An T. Nguyen, Matthew A. Albrecht, Ottmar V. Lipp, Welber Marinovic

**Author notes:** **Corresponding authors** School of Psychology, Building 401, Bentley 6102, Perth, WA, Australia, **E-mail:**.

## Abstract

Motor actions can be suppressed with varying degrees of success, but this variability is not captured in many experiments where responses are represented in binary (response vs. no-response). Although the Stop/No-go P300 (an enhanced frontocentral positivity in the event-related potential (ERP) peaking around 300 ms after a Stop/No-go stimulus, compared to Go trials) has been implicated as a measure of inhibitory-control, it is unclear how the range of motor outputs relates to the P300. We examined the nature of this association in two experiments using an Anticipatory Timing and a Go/No-go Task. Force, response onset time, and the P300 were measured.

In both experiments, our results showed that trial-by-trial P300 amplitude on Failed Inhibitions were continuously related to Force, where higher response forces (reflecting a greater degree of error) were associated with smaller P300 amplitudes. Compared to Successful Inhibitions, P300 amplitude was reduced and ERP onset latencies were delayed on Failed Inhibitions. Although the binary categorisation of inhibition-success (Successful vs. Failed) accounted for more variance in the data compared to force, it misses a reliable linear relationship that can be captured by continuous measures of motor output. Overall, the results provide strong evidence that the engagement of inhibitory-control varies on a continuum from trial-to-trial and that this engagement is reflected by the P300. We present an activation-to-threshold model of inhibitory-control to explain our results, which offers a new conceptual framework for describing the implementation of inhibitory-control in highly prepared motor responses. Our results also highlight the importance of studying the spectrum of motor outputs and the need for future models of inhibitory-control to account for motor output.

## 1. Introduction

Elite athletes in sports such as tennis and baseball, have little time to prepare and execute successful actions (e.g. hitting a fast pitch). To deal with these high temporal demands humans rely on advanced motor planning (de Rugy, Loeb, & Carroll, 2012; Gray, 2002; Lacquaniti & Maioli, 1989; Marinovic, Tresilian, Chapple, Riek, & Carroll, 2017; Reuter, Marinovic, Welsh, & Carroll, 2019; Senot, Zago, Lacquaniti, & McIntyre, 2005). Although this strategy increases the chances of success when our predictions are correct, it becomes challenging to change or inhibit these highly prepared actions when they become undesirable (e.g. not hitting a ball that is going ‘out’, checked swings in baseball). Current psychophysiological models conceptualise the outcome of the inhibitory process as a binary event in which responses are either suppressed or not (e.g. Horse Race Model; Logan & Cowan, 1984; Logan, Van Zandt, Verbruggen, & Wagenmakers, 2014; Verbruggen et al., 2019), reflecting the binary measures (e.g. key-presses) often used to record responses. Motor actions, however, can be inhibited with varying degrees of success (de Jong, Coles, Logan, & Gratton, 1990; Marinovic, Plooy, & Tresilian, 2009; Marinovic, Reid, Plooy, Riek, & Tresilian, 2010; Marinovic, Reid, Plooy, Tresilian, & Riek, 2010; Nguyen, Moyle, & Fox, 2016; Scheffers, Coles, Bernstein, Gehring, & Donchin, 1996), ranging from no output on Successful Inhibitions, to slight twitches and larger movements on Partial and Failed Inhibitions, respectively. In this study, we examined how the range of motor outputs relates to electrophysiological measures of inhibitory-control.

A common approach to studying inhibitory-control is to assign responses to discrete categories (e.g. Correct or Incorrect Response, Successful or Failed Inhibition) with little regard to the fact that these responses are likely to be distributed in a continuum. This assumes that all responses within a category are equivalent, which is not necessarily the case. Behavioural experiments have shown that the timing of the response on the correct limb is interfered by the additional presence of a partial response on the incorrect limb, indicating that not all correct trials are alike (Coles, Gratton, Bashore, Eriksen, & Donchin, 1985; Eriksen, Coles, Morris, & O’hara, 1985). Similarly, electrophysiological studies have shown that partial errors elicit reduced error monitoring signals compared to complete errors (Meckler, Carbonnell, Ramdani, & Hasbroucq, 2017; Vidal, Hasbroucq, Grapperon, & Bonnet, 2000). This highlights that not all errors are comparable and suggests that differences in the brain’s responses may be related to motor output and vice-versa.

In inhibitory-control tasks, the Stop/No-go P300 event-related potential (ERP) component has been associated with the inhibitory-control process (Pires, Leitão, Guerrini, & Simões, 2014). It is typically enhanced on Inhibition compared to Go or non-inhibition trials (de Jong et al., 1990; Dimoska, Johnstone, & Barry, 2006; Randall & Smith, 2011; Smith, Johnstone, & Barry, 2008), and has been shown to be earlier and/or larger on Successful compared to Failed Inhibitions (Carrillo-de-la-Peña, Bonilla, & González-Villar, 2019; de Jong et al., 1990; Dimoska & Johnstone, 2008; González-Villar, Bonilla, & Carrillo-de-la-Peña, 2016; Greenhouse & Wessel, 2013; Kok, Ramautar, De Ruiter, Band, & Ridderinkhof, 2004; Liotti, Pliszka, Perez, Kothmann, & Woldorff, 2005; Overtoom et al., 2002; Schmajuk, Liotti, Busse, & Woldorff, 2006; Wessel & Aron, 2015). Interestingly, we have discovered that partial failures to inhibit (erroneously initiated responses that were cancelled before completion) also resulted in significant reductions in P300 amplitude (Nguyen et al., 2016). Motor output related reductions in P300 have also been observed in comparisons between Count vs. Press Go/No-go Tasks and in motor-execution tasks (Novembre et al., 2018; Smith et al., 2008). Collectively, there is some evidence suggesting a possible relationship between P300 and motor output, but the nature of this association remains to be examined.

In the current study, we took advantage of the temporal control provided by the Anticipatory Timing Task to gain insight into the nature of the relationship between P300 and motor output. In the Anticipatory Timing Task, participants anticipated and synchronised their responses to a predictable event, while inhibiting their responses to a Stop-signal. It is similar to the Stop-Signal Task in that it requires participants to cancel their response during the late phases of response preparation and execution. However, it differs in its anticipatory design, absence of a reaction to a separate Go stimulus, and the use of a fixed Stop-signal time. This design allows us to eliminate any differences in stimulus-presentation across inhibition trials and across participants, which may complicate the interpretation of modulations in the ERP. We also conducted an additional experiment using a Go/No-go task to examine if this relationship can be observed in other inhibitory-control tasks. If inhibitory-control is reflected by the P300, we predicted that it should be modulated by the success of inhibition, producing enhanced and earlier P300s on Successful compared to Failed Inhibitions. If the inhibitory-control process can vary continuously in its engagement from trial-to-trial, we expected that P300 amplitude would be negatively related to the force of responses on Failed Inhibitions.

## 2. Methods

### 2.1. Participants

For each experiment, we recruited 28 participants consisting of university students and volunteers. Three participants were excluded from each study due to missing data or excessive EEG noise and artifacts (Exp. 1, final N = 25, M(SD) age = 21.16 (2.94) years, range = 18-27 years; Exp. 2, final N = 25, M(SD) = 21.56(3.31) years, range = 18-30 years). The study was approved by the human research ethics committee of Curtin University (Approval code: HRE2016-0200) and all participants provided written informed consent before completing the experiment. All participants also reported being right-handed, having normal or corrected vision, and not having any known or diagnosed neurological conditions.

### 2.2. Experiment 1: Anticipatory Timing Task

Participants were instructed to synchronise their response with the intercept of the yellow rectangle and the green target, unless a Stop-signal was presented (Fig. 1). Participants responded by performing brief isometric wrist-extensions with their right arm, which was secured inside a custom-built wrist-device (de Rugy et al., 2012). The anticipatory design allows us to overcome some of the methodological challenges associated with examining ERPs in Stop-Signal Tasks. In the Stop-Signal Task, the short time gap between the presentation of Go and Stop-signal stimuli is known to produce overlapping neural activity. Additionally, the relative timing of the Stop-signal also varies from trial-to-trial. These varied and overlapping activations can complicate the interpretation of changes in the ERP. By using the Anticipatory Timing Task, which does not present a Go stimulus that participants must react to, we were able to minimise effects associated with orienting and processing the Go stimulus, and variability in Stop-signal timings was minimised by using a fixed Stop-signal timing. To ensure the task was difficult enough to elicit Failed Inhibitions, the Stop-signal was always presented 175 ms before the intercept, which has been shown to translate to ~50 % Successful Inhibitions across participants based on previous findings using the Anticipatory Timing Task (Marinovic et al., 2009; Marinovic, Reid, Plooy, Riek, et al., 2010; Marinovic, Reid, Plooy, Tresilian, et al., 2010). To match the presentation of stimuli between Go and Stop-trials, an ‘Ignore Go’ condition was included where participants were instructed to ignore an upcoming Stop-signal.

**Figure 1.**
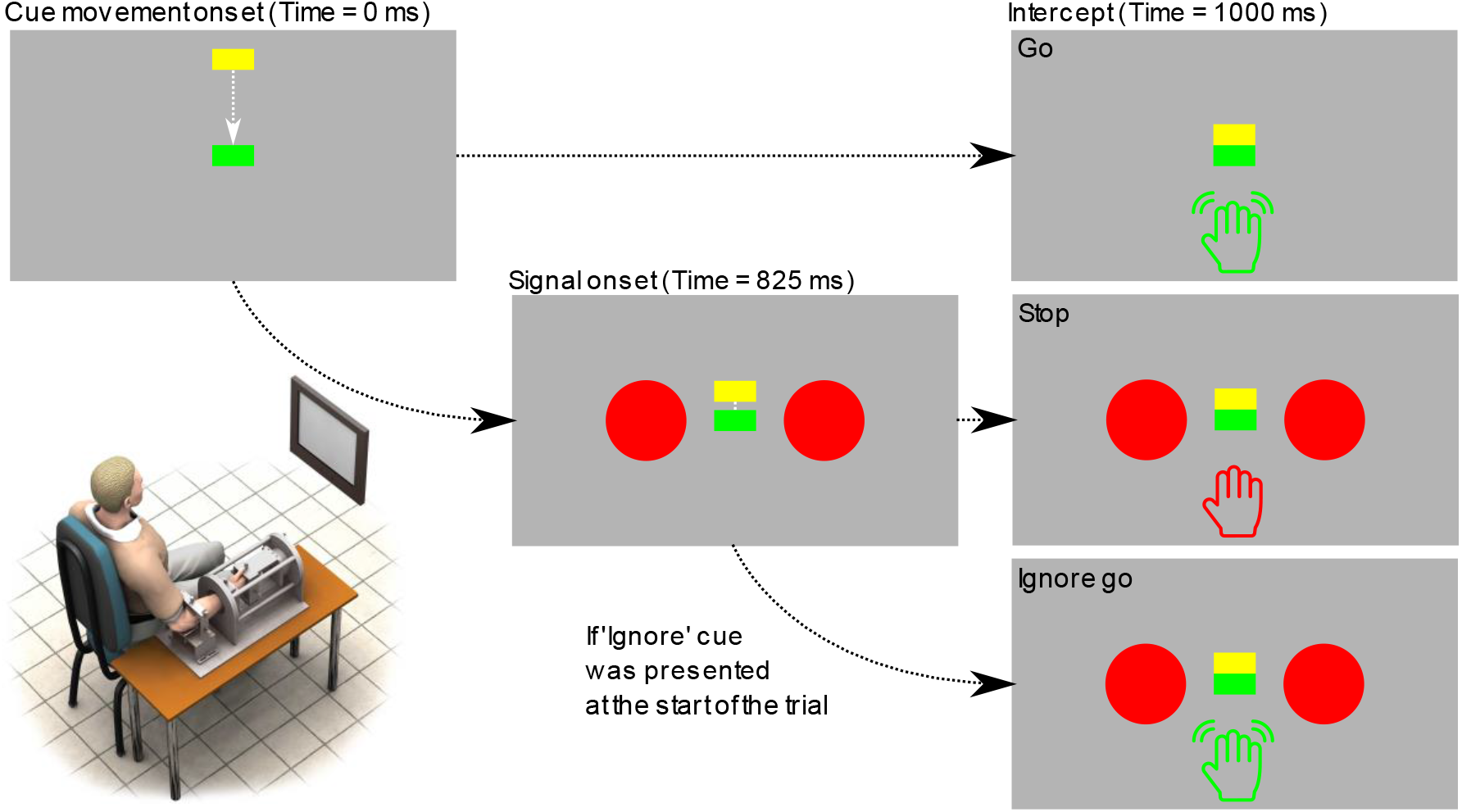
Diagram of the Anticipatory Timing Task. Participants were instructed to respond when the yellow rectangle reached the green target (Go trial) and to withhold their responses if a Stop-signal was presented (Stop trial). On Ignore Go trials, participants were presented with a cue at the beginning of the trial, instructing participants to respond to the trial despite the presentation of the Stop-signal. Participants responded by performing brief isometric wrist-extensions.

The task was presented on a 24-inch LCD monitor (1920 x 1080 resolution, 120 Hz) using MATLAB 2015b and Psychtoolbox version 3.0.11 (Brainard, 1997; Kleiner et al., 2013; Pelli, 1997). At the start of each trial, the text “*Relax*” was presented for 1000 ms. On Ignore Go trials, the text “*Ignore”* was presented for 1500ms, accompanied by an audible tone. Yellow and green rectangles (100 x 50 pixels) were then presented at the top and middle of the display for 500 ms (Fig. 1). The yellow rectangle descended, intercepting with the green target after 1000 ms. On Stop and Ignore Go trials, two red circles (300-pixels diameter each) were presented on the left and right of the green box, 175 ms *before* the intercept (i.e. 825 ms after the yellow rectangle starts moving). Visual feedback about temporal accuracy and force was presented for 500ms after the intercept for 1000 ms.

Movement onset and peak force (in Newtons) were the measures of response time and force. We measured responses using a custom-built wrist-device with a six-degree of freedom force/torque sensor (JR3 45E15AI63-A 400N60S, Woodland, CA) digitised using a National Instruments data acquisition device (USB-6229 BNC multifunctional DAQ). Although the sensors are capable of six degrees of freedom, only movements in the horizontal plane were required for this experiment. We calculated movement onset from the tangential speed time-series derived from the torque data using the algorithm recommended by Teasdale and colleagues (1993).

Participants completed two practice and three experimental blocks (10, 50 and 3 × 100 trials, respectively). The first practice block consisted of all Go trials, while all other blocks consisted of 40 % Go, 40 % Stop and 20 % Ignore Go trials, presented in a pseudo-random order. While the global percentage of response trials was 60 %, the conditional probabilities of Go and Stop-trials were equal, minimising the novelty-related effects which are known to influence the P300 and may confound inhibition-related effects (Debener, Makeig, Delorme, & Engel, 2005; Dimoska & Johnstone, 2008; Donkers & van Boxtel, 2004; Ramautar, Kok, & Ridderinkhof, 2004). Participants were instructed to time the onset of their responses such that it coincided with the intercept, and to produce moderate forces of at least 20 N. As we wanted participants to focus on the timing of their responses, we only defined a moderate minimum force level and did not specify any additional constraints on force. Correct Go trials were identified as those with responses starting within −50 to 30 ms from the intercept, but participants were not explicitly aware of this criterion. A shorter post-intercept time was selected to discourage participants from delaying their responses in anticipation of the Stop-signal. Visual feedback was presented at the end of each trial (e.g. “Good”, “Too Early”, “Too Soft”). Numerical force values were only presented as feedback during practice blocks. On Go trials, participants were informed if responses were too early or late (−50 < RT > 30 ms), or too soft (< 20 N). Successful Stop-trials were defined as Stop-trials without a detectable response. Failed Stops were defined as Stop-trials with a detectable response within −175 ms to 500 ms from the intercept. Responses initiated outside of this interval were considered out-of-bounds and were excluded from further analyses. The same feedback was presented for Failed Stop and out-of-bound trials, informing participants of an incorrect response.

### 2.3. Experiment 2: Go/No-Go Task

Participants were instructed to quickly respond to Go stimuli while withholding responses to No-go stimuli (Fig. 2). At the start of each trial, the text “Relax” was presented for 1000 ms, followed by a warning cue (hollow white circle, 300-pixel diameter) for a randomly varied duration between 1000 to 1200 ms. A Go or No-go stimulus (blue or yellow circle, 300-pixel diameter, counter-balanced across blocks and participants) was then presented for 500 ms, followed by visual feedback for 1000 ms.

**Figure 2.**
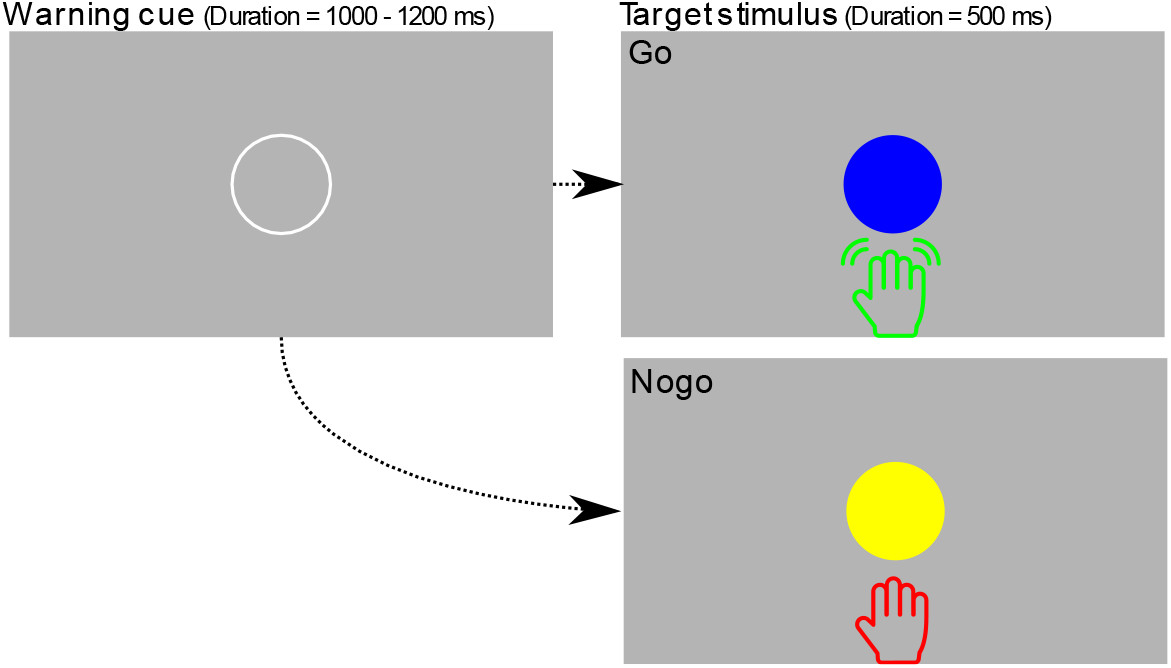
Diagram of the Go/No-go Task. Participants were instructed to quickly respond to Go stimuli (within 200 ms) while withholding responses to No-go stimuli. The coloured stimuli were counter-balanced across blocks for each participant.

Participants completed two practice and four experimental blocks (10, 24 and 4 × 80 trials, respectively). The first practice block consisted of all Go trials while all other blocks consisted of 50 % Go and No-go trials, presented in a pseudo-random order. An equiprobable task was used to match the design of the first experiment. Participants were instructed to initiate their response to Go stimuli within 200 ms to encourage the development of a pre-potent response, and to produce moderate forces of at least 20 N. Correct Go trials were identified as those with responses starting within 200 ms of the stimulus. Visual feedback was presented at the end of each trial. On Go trials, participants were informed if their responses were too early or late (0 < RT > 200 ms), or too soft (< 20 N). On No-go trials, Successful No-go trials were defined as trials without a detectable response. Failed No-go trials were defined as trials with a detectable response within 0 to 500 ms after stimulus onset. No-go trials with responses initiated outside of this window were considered out-of-bounds and were excluded from further analyses. To encourage fast responses, we awarded participants with 3 points for each Correct Go response and 1 point for each Successful Inhibition. No points were deducted for incorrect responses and the points were reset after each block.

### 2.4. EEG Acquisition and Pre-Processing

EEG data were recorded continuously for the duration of the experimental blocks. Data were acquired using a Biosemi ActiveTwo EEG system and ActiView (ver. 7.07) at a sampling rate of 1024 Hz with a DC – 100 Hz online filter. Data were recorded from 64 scalp electrodes arranged according to the 10-20 system with additional electrodes placed adjacent to the outer canthi of both eyes and on the left infraorbital region. For online referencing, the Biosemi EEG system uses active electrodes with common mode sense (CMS) and driven right leg (DRL) electrodes providing a reference relative to the amplifier reference voltage.

EEG data were processed offline in MATLAB 2018a using EEGLAB (Delorme & Makeig, 2004), AMICA (Palmer, Kreutz-Delgado, & Makeig, 2011), SASICA (Chaumon, Bishop, & Busch, 2015), and ERPLAB (Lopez-Calderon & Luck, 2014) plugins. The data were re-referenced to the average of the 64 scalp electrodes, down-sampled to 256 Hz and filtered from 0.1 – 40 Hz (−6 dB) using separate low- and high-pass Kaiser filters with transition widths 0.2 and 2 Hz, a maximum passband deviation of .001, and filter orders of 4638 and 466.

Epochs were extracted for Correct (Ignore) Go, Successful and Failed Inhibitions, time-locked to the presentation of the Stop-signal/No-go/Go stimulus. Epochs spanned from −1100 to 1200 ms, and baseline amplitudes were corrected to the 100 ms interval preceding the time-locked stimulus. To correct for blinks, horizontal saccades and other artifacts, Independent Component Analysis was conducted and independent components (ICs) containing artifacts were manually identified with the guidance of SASICA and removed (Exp. 1: M(SD) = 13.4(7.3) ICs; Exp. 2: M(SD) = 12.6(7.2) ICs). Trials containing voltages exceeding ±100 μV were excluded (Exp. 1: M(SD) = 5(6.28) trials, max = 16 trials; Exp. 2: M(SD) = 3.57(2.82) trials, max = 8 trials). For experiment 1, after trial rejection, the average number of trials retained for Correct Ignore Go, Successful and Failed Stops were 27.88(8.02), 67.6 (18.97) and 51.52(18.97), respectively. For experiment 2, the average number of trials retained for Correct Go, Successful and Failed No-go conditions were 78.36 (31.90), 112.52 (25.66), and 39.80 (22.14), respectively. A Surface Laplacian filter was applied using algorithms described in Perrin and colleagues (1989) (smoothing factor = 1e^-5^, order of Legendre polynomial = 10) to reduce volume conduction effects in EEG sensor space, resulting in a μV/mm^2^ voltage scale.

### 2.5. Measuring ERP Amplitude and Onset Latency

To emphasise inhibition-related activity, we focused our analyses on ‘Inhibition minus Go’ difference waveforms (Stop minus Correct Ignore Go, No-go minus Correct Go). To retain trial-level information, the averaged Correct (Ignore) Go ERP was subtracted from each inhibition trial waveform. To compensate for increased noise associated with trial-level analyses, we used an averaged region of frontocentral channels (FC1, FCz, FC2, C1, Cz, C2) for measurement. Inspection of our data showed that P300 was maximal in the frontocentral region (Fig. 3).

**Figure 3.**
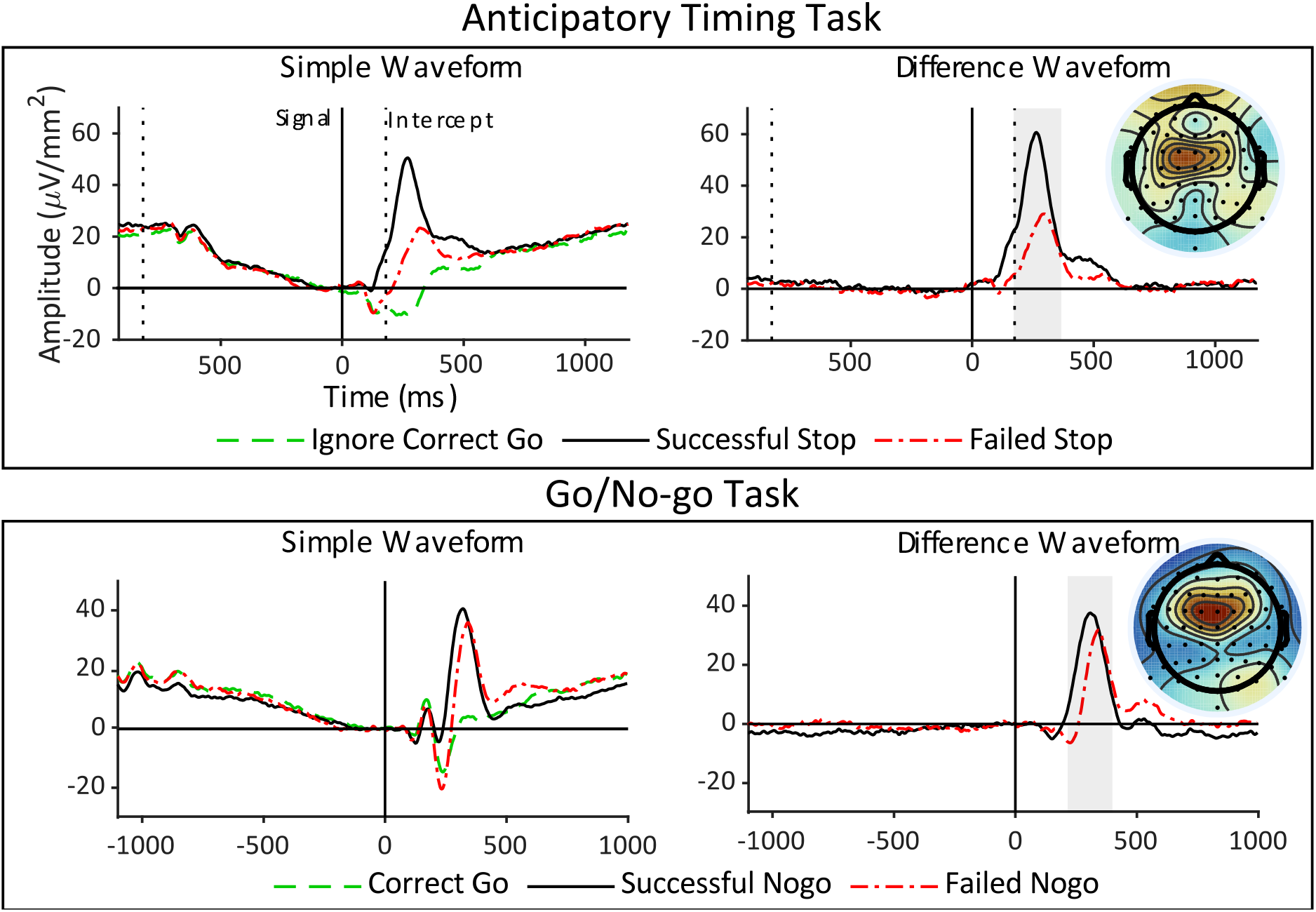
Grand-averaged waveforms of an averaged frontocentral region (FC1, FCz, FC2, C1, Cz, C2). Difference waveforms reflect ‘Inhibition - Correct (Ignore) Go’ differences. Shaded areas show measurement intervals for P300. Scalp maps reflect mean amplitudes over the measurement interval for Successful Inhibitions.

We measured P300 mean amplitude over a 200 ms interval centred on the positive peak (200-500 ms post-stimulus) of the grand-averaged Successful Inhibition difference waveform (Fig. 3). With respect to latency, we measured the onset of ‘Inhibition minus Go’ difference waveforms using the ‘Median Rule’ described in Letham and Raij (2011). This method compares each point in the time-series with a threshold based on the median and the interquartile range of baseline activity. Due to difficulty detecting onsets at the trial-level, onset latencies were measured on individual-averaged difference ERPs on Successful and Failed Inhibitions. To reduce the chance of selecting transient noise spikes, we required 10 consecutive time-points to exceed threshold by 100 ms post-stimulus and onset latencies refer to the first point in this sequence. For experiments 1 and 2 respectively, we excluded 5 and 1 participants where onset latency could not be identified or who showed floor effects in both Successful and Failed Inhibitions. These participants were only excluded for the onset latency analysis.

### 2.6. Statistical Analysis

Three sets of statistical analyses were conducted using linear mixed models in R, with the ‘lmer’ function from the ‘lmer est’ package (Kuznetsova, Brockhoff, & Christensen, 2017). Participant intercepts were modelled as a random effect in all analyses. We used the ‘emmeans’ function from the ‘emmeans’ package (Lenth, Sigmann, Love, Buerkner, & Herve, 2019) to present the results of categorical comparisons as *t*-ratios with degrees-of-freedom estimated using the Kenward-Roger method. e used the ‘ano a’ function to present the results of trial-le el analyses as F alues using the atterthwaite’s approximation method (Satterthwaite, 1941). We also used the ‘summ’ function in the ‘jtools’ package (Long, 2019) and the ‘r2beta’ function from the ‘r2glmm’ package to calculate the estimated regression-weights (*β*) and effect size as Partial *R^2^* (*R_p_*^2^) (Jaeger, 2017).

Firstly, we compared force and response time between Go and Failed Inhibitions, and we examined the association between force and response time on Failed Inhibitions. The former was done by modelling force and response time at the participant-level with trial-category (Go, Failed Inhibition) as a fixed-effect. The latter was done by modelling response time at the trial-level with force as a fixed effect. Response time and force were scaled by participant in this analysis to reduce scale differences between the two variables.

Secondly, we examined whether P300 was modulated by the success of inhibitory-control. We modelled P300 difference mean amplitude and ERP difference onset latencies at the participant-level with binary inhibition-success (Successful, Failed) as a fixed effect. Using the same approach, we also compared ERP difference onset latency with response time on average Go trials.

Thirdly, we examined the association between P300 amplitude and motor output on Failed Inhibitions, and we examined how force and binary inhibition-success compare as predictors of P300 amplitude. The former was done by modelling P300 difference mean amplitude on Failed Inhibitions at the trial-level with force as a fixed-effect. The latter was done by modelling P300 difference mean amplitude on all inhibition trials with force and inhibition-success (Successful, Failed) as fixed effects. Force was scaled by participant in this analysis to focus on relative changes in force within-individuals.

## 3. Results

### 3.1 Force, Response Time and Inhibition Accuracy

Participants successfully inhibited their responses on 56.33(15.81) % and 70.33(16.04) % of trials in the Anticipatory Timing and Go/No-go Tasks, respectively. Unsuccessfully inhibited responses were significantly less forceful than the average Go response, suggesting that some degree of movement suppression was present (Fig. 4; Exp. 1: 17.08(12.99) vs. 32.07(13.4) N, *t-ratio* (24) = 10.94, *p* < .001; Exp. 2: 9.99(11.71) vs. 37.32(13.1) N, *t-ratio*(24) = 12.51, *p* < .001). With respect to timing, unsuccessfully inhibited responses were initiated significantly earlier than the average Go response, suggesting that these responses may have been more highly prepared compared to the average response and were therefore more difficult to suppress (Fig. 4; Exp. 1: −43.32 (54.50) vs. 1.19 (63.16) ms relative to the intercept, *t-ratio*(24) = 8.91*, p* < .001; Exp. 2: 154.14 (64.89) vs. 209.4 (62) ms relative to stimulus-onset, *t-ratio*(24) = 16.14, *p* < .001). When examining the association between response time and force for Failed Inhibitions, we found that response time was negatively related to force in the Anticipatory Timing Task, indicating that earlier responses tended to be more forceful (*F*(1,1286) = 27.60, *p* < .001, *R*^2^_*p*_ = 0.02, *β* = −0.14). However, this association was not statistically significant in the Go/No-go Task (*F*(1,993) = 2.20, *p* = 0.139, *R*^2^_*p*_ = 0.002, *β* = −0.05).

**Figure 4.**
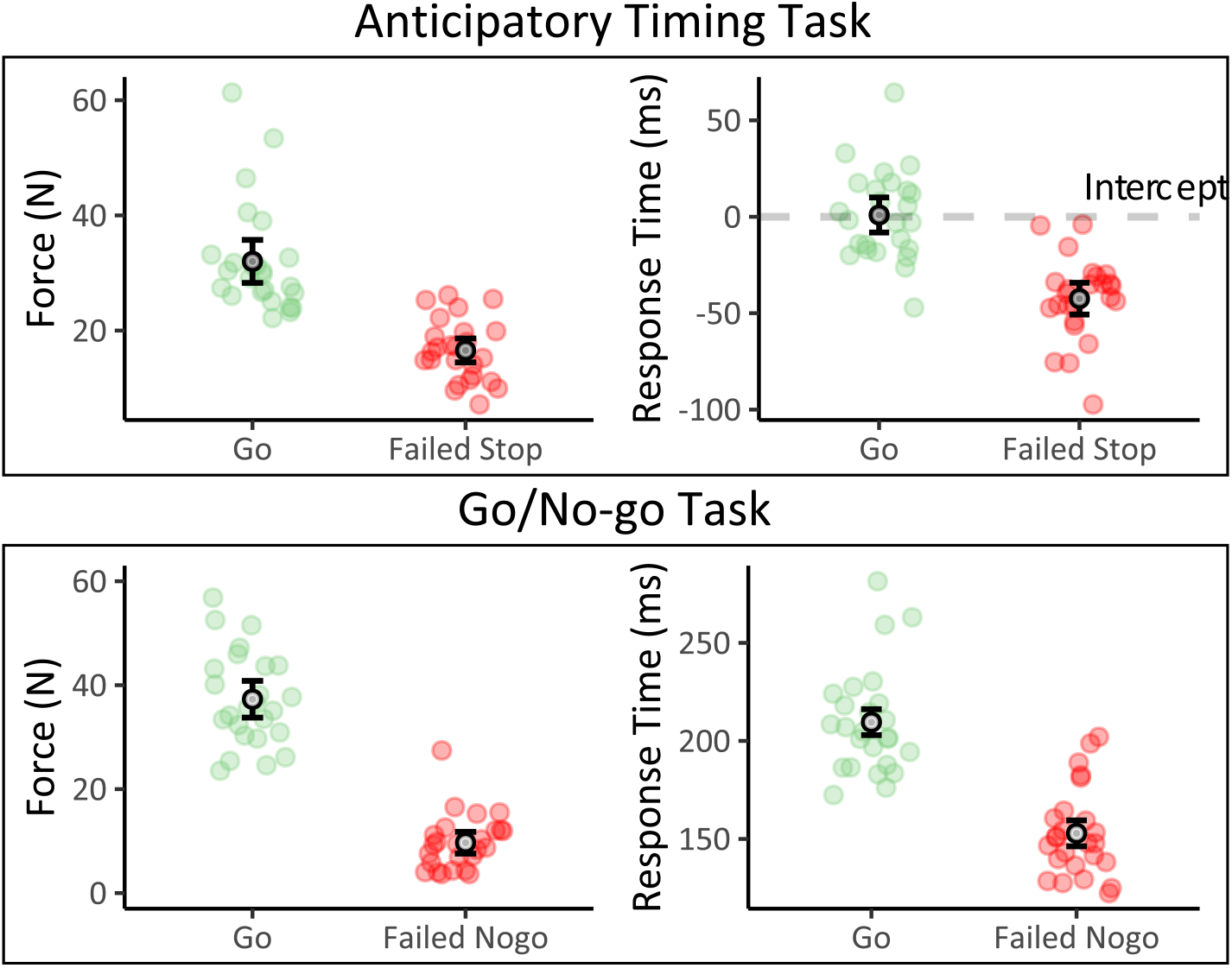
Force and response time plots of individual- and grand-averages with 95% confidence interval error-bars. Response times in the Anticipatory Timing Task are relative to the intercept shown by the dashed grey line. Response times in the Go/No-go task are relative to the onset of the Go and No-go stimuli.

### 3.2. P300 Difference Mean Amplitude and ERP Difference Onset Latency

P300 difference mean amplitude was significantly reduced on Failed compared to Successful Inhibitions (Fig. 5; Exp. 1: 18.66(10.43) s. 40.39(15.56) μV/mm^2^, *t-ratio* (24) = 8.89, *p* < .001; Exp. 2: 13.43(11.55) s. 24.57(14.30) μV/mm^2^, *t-ratio* (24) = 9.669, *p* < .001). With respect to latency, ERP difference onset was significantly delayed for Failed compared to Successful Inhibitions (Fig. 5; Exp. 1: 191.80(60.93) vs. 135.35(39.94) ms relative to Stop-signal, *t-ratio* (19) = 4.23, *p* < .001; Exp. 2: 252.44(49.04) vs. 219.40(24.65) ms relative to no-go stimulus, *t-ratio* (23) = 2.70, *p* = .013).

**Figure 5.**
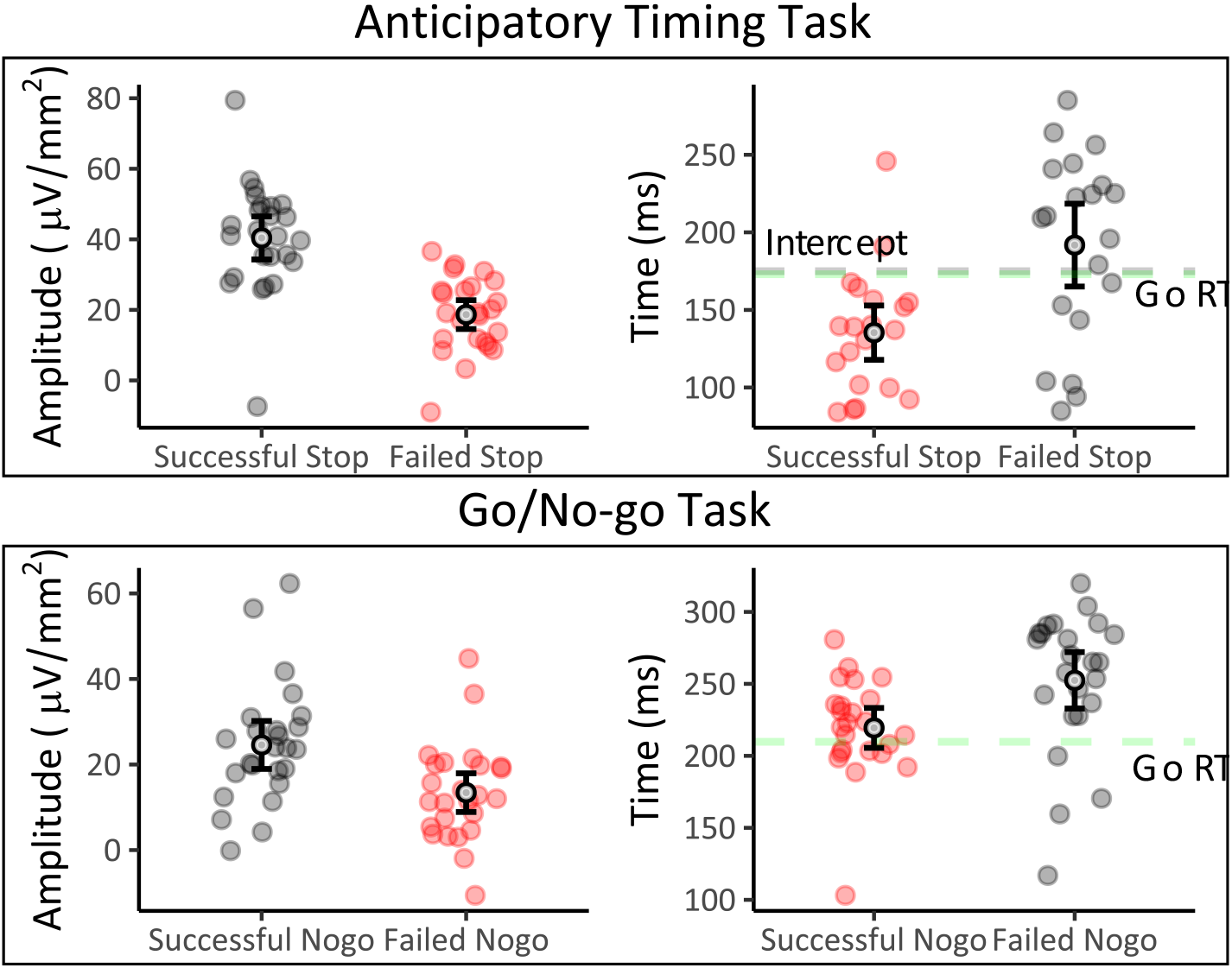
P300 Difference Mean Amplitude and ERP Difference Onset Latency plots of individual- and grand-averages with 95% confidence interval error-bars. Onset times are relative to the onset of the Stop/No-go stimulus. Dashed grey and green lines represent the intercept and average Go response time, respectively.

When comparing ERP difference onset latency with response time, we found that the onset of the ERP difference occurred before the average response on Go trials in the Anticipatory timing task, indicating that inhibitory-control was engaged before the response was initiated (−39.65(39.94) vs. −1.99(24.93) ms relative to the intercept, *t-ratio* (19) = 3.58, *p* = .002). However, ERP difference onset latency did not precede the average response time on Go trials in the Go/No-go task. Rather, our results indicated that they coincided temporally (215.92(32.45) vs. 209.78(28.29) ms relative to stimulus onset, *t-ratio* (23) = 1.40, *p* = .176).

### 3.3. Association between P300 Difference Mean Amplitude and Motor Output and Inhibition-Success on Failed Inhibitions

Force was a significant negative predictor of P300 amplitude on Failed Inhibitions, where greater forces (reflecting a higher degree of error) were associated with reduced amplitudes (Exp. 1: *F*(1,1262.5) = 32.06, *p* < 0.001, *R*^2^_*p*_ = .025, *β* = −3.24; Exp. 2: *F*(1,968.59) = 49.31, *p* < 0.001, *R*^2^_*p*_ = .048, *β* = −4.47). When modelling P300 amplitude on all inhibition trials with force and binary inhibition-success, both variables were significant negative predictors of P300 amplitude. However, the proportion of variance explained by force — a continuous predictor— was smaller compared to inhibition-success (Exp. 1: Force, *F*(1,2947.1) = 43.61, *p* < 0.001, *R*^2^_*p*_ = .015, *β* = −3.47; Binary Inhibition-Success, *F*(1,2954.9) = 210.74, *p* < 0.001, *R*^2^_*p*_ = .067, *β* = −16.17; Exp. 2: Force, *F*(1,3776.1) = 47.32, *p* < 0.001, *R*^2^_*p*_ = .012, *β* = −2.79; Binary Inhibition-Success, *F*(1,3785.6) = 53.79, *p* < 0.001, *R*^2^_*p*_ = .014, *β* = - 7.15).

## 4. Discussion

In this study, we investigated the nature of the relationship between motor output and P300, to gain further insight about its implicated role as an electrophysiological measure of inhibitory-control. Using the Anticipatory Timing and the Go/No-go tasks, we measured the force and the timing of responses, and the frontocentral P300 elicited by inhibition stimuli. In our analyses, we characterised the behavioural and electrophysiological response to Failed Inhibitions and examined the relationship between P300 and motor output. If inhibitory-control is reflected by the P300, it should be modulated by the success of inhibition. As such, we expected enhanced and earlier P300s on Successful compared to Failed Inhibitions. We also expected that P300 amplitude on Failed Inhibitions to be negatively related to force, suggesting a continuous relationship between the engagement of inhibitory-control and the degree of inhibition failure.

In regards to behaviour, unsuccessfully inhibited responses were less forceful than the average Go response, highlighting that there was some degree of response suppression on Failed Inhibitions which is consistent with previous studies examining response force (Ko, Alsford, & Miller, 2012; Scheffers et al., 1996). Response times were also shorter for Failed Inhibitions, suggesting that they were more highly prepared compared to the average Go response, possibly contributing to the failure in completely inhibiting these responses. With respect to electrophysiology, our results were in line with our predictions. P300 was both reduced and the onset of ERP differences were delayed on Failed compared to Successful Inhibitions. Our linear modelling results revealed a significant negative relationship between amplitude and force, where greater force (reflecting a higher degree of error) was associated with smaller amplitudes. The relationship between motor output and P300 is consistent with our previous work on partial inhibitions and recent work examining motor output (Nguyen et al., 2016; Novembre et al., 2018).

Supporting the P300 as an electrophysiological measure of inhibitory-control, our latency analysis showed that the onset of differences in activation between Successful Inhibition and Go trials significantly preceded the average timing of Go responses in the Anticipatory Timing Task, indicating that the inhibitory-control process was engaged prior to the execution of the response. While our onset times are relatively short compared with the P300 onsets commonly observed in the Stop-Signal Tasks, they are in line with the timing of inhibitory effects seen in studies using transcranial magnetic stimulation (TMS) (Badry et al., 2009; Wildenberg, Burle, Vidal, Molen, & Ridderinkhof, 2009). However, onset latencies in the Go/No-go Task did not precede the timing of average Go responses. Rather, our results suggested that they coincided temporally. Given the temporal proximity between the onset of the stimulus and the response (< 210 ms), we cannot discount the possibility that modulations in P300 may also reflect the post-response evaluative processing, an idea that has been suggested in previous Go/No-go studies (Bruin, Wijers, & van Staveren, 2001; Huster, Enriquez-Geppert, Lavallee, Falkenstein, & Herrmann, 2013; Nguyen et al., 2016).

In addition to discrepancies in onset latency, differences in P300 amplitude between Successful and Failed Inhibitions were also noticeably smaller in the Go/No-go compared to the Anticipatory Timing Task. These discrepancies may be related to the influence of anticipatory- and reaction-based task designs on response preparation and inhibitory-control. In the Anticipatory Timing Task, the anticipatory nature of responding to a known event facilitates advanced response-preparation, thereby increasing the amount of inhibitory-control required to suppress highly prepared responses. Comparatively, the presentation of the Go stimulus is relatively uncertain in the Go/No-go task which may reduce preparation, thereby decreasing the amount of control required. Similarly, previous comparisons between the Stop-Signal and Go/No-go tasks have reported larger P300 amplitudes in the Stop-Signal task, where the Go stimulus precedes the Inhibitory stimulus, allowing for the advance preparation and execution of the response (Enriquez-Geppert, Konrad, Pantev and Huster, 2010). Differences between the Go/No-go and the Anticipatory Timing Task could also point to the engagement of distinct but related neural circuits, as has been shown in a meta-analysis comparing Stop-signal and the Go/No-go task (Swick, Ashley, & Turken, 2011).

With respect to the inhibition-success effect on P300 amplitude, differences in reward feedback may have also contributed to task discrepancies, as the awarding of points in the Go/No-go Task was another point of difference, introduced to encourage pre-potent responding in the equiprobable task. In a previous study exploring the role of motivation on inhibition-success effects, it was demonstrated that biasing rewards towards correct Go responses reduced P3 differences between Successful and Failed Inhibitions which is in line with our observed results (Greenhouse & Wessel, 2013). It was suggested that this modulation may reflect underlying attention-driven processes, but it is not clear whether the process itself is response-inhibition. Collectively, these discrepancies highlight the possibility that the P300 may reflect a combination of inhibitory-control and evaluative processing, as well as the impact of task-design on inhibitory-control and on factors such as motor-preparation and motivation. However, despite these discrepancies, the pattern of inhibition and force-related effects on P300 was maintained across experiments, demonstrating the generalisability of our results.

Lastly, we examined the contribution of force on P300 and how it compares to the binary categorisation of inhibition-success by modelling P300 amplitude across all Inhibition trials. Although both variables were statistically significant predictors, the binary categorisation of inhibition trials accounted for a larger proportion of variance. This result indicates that the difference in P300 amplitude between Successful and Failed Inhibitions is greater than changes across the spectrum of Failed Inhibitions. Given that motor output on Failed Inhibitions was associated with varying levels of inhibitory-control engagement reflected by the P300, Successful Inhibitions may also be associated with a range of engagement levels. We suggest that the binary effect arises from the grouping of higher and lower levels of the inhibitory-control engagement continuum, but this does not necessarily mean that the inhibitory-control process is all-or-none. The larger effect of binary categorisation compared to force can simply indicate that there is a larger range of inhibitory-control engagement levels that are associated with Successful Inhibitions that are behaviourally unobserved. Although the binary categorisation of Successful and Failed Inhibitions accounts for a larger proportion of the variance, it misses a reliable linear relationship between P300 and motor response that is only captured in continuous measurements of motor output. This association is a crucial piece of evidence which provides important information about the functional significance of the P300 and its role in inhibitory-control.

### 4.1. Activation-to-Threshold Model of Inhibitory-Control

To illustrate our conceptualisation, we present an activation-to-threshold model of inhibitory-control for the Anticipatory Timing Task (Fig. 6). The model consists of opposing preparatory and inhibitory processes, where these processes can vary freely on a continuum from trial-to-trial as expectancies about the timing of movement onset and the appearance of a Stop-signal fluctuate. The model predicts the presence or absence of a response, and its force and timing of response onset are based on the combined (net) activation of these competing processes. The model was derived from previous studies using the Anticipatory Timing Task to explain vigour and response time effects during loud acoustic stimulation (Marinovic, de Rugy, Lipp, & Tresilian, 2013; Marinovic, Tresilian, de Rugy, Sidhu, & Riek, 2014; Tresilian & Plooy, 2006). More recently, similar activation-to-threshold concepts have also been described in the context of inhibitory-control (MacDonald, McMorland, Stinear, Coxon, & Byblow, 2017).

**Figure 6.**
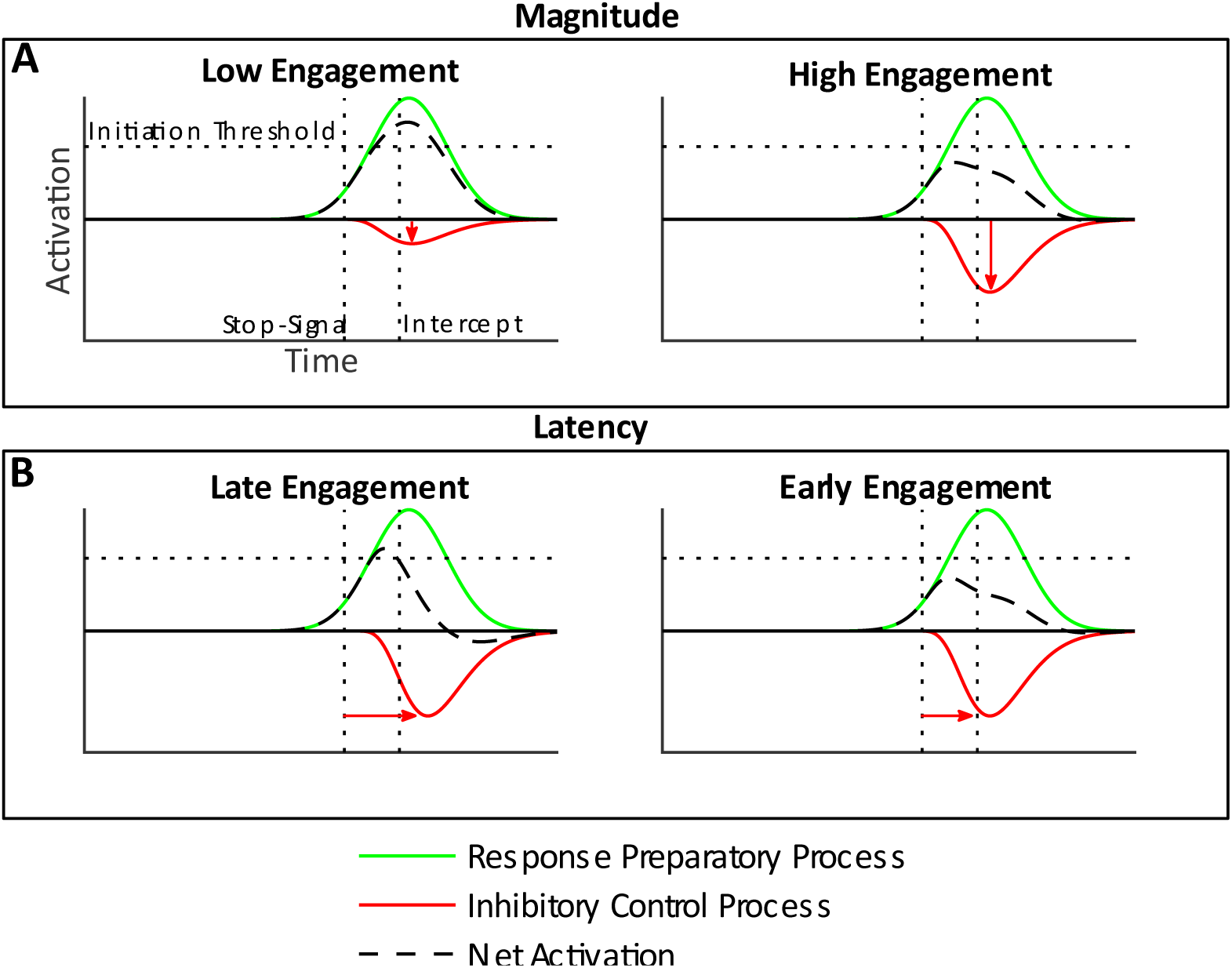
A graphical depiction of the Activation-to-Threshold Model of Inhibitory-Control for the Anticipatory Timing Task. The model shows the activity of opposing preparatory (green) and inhibitory processes (red), and their combined signal (net activation, dashed black). A response is initiated if net activation crosses the initiation threshold, and its timing and force are determined by the point at and the extent to which the threshold is crossed. For a given level of response preparation, the model shows how differences in the magnitude and latency in the engagement of the inhibitory-control process can result in Failed and Successful Inhibitions.

We modelled that response activation would increase close to the intercept as the participant prepares to respond, increasing net activation. When the Stop-signal is presented, the inhibitory-control process is rapidly engaged (similar to models describing the Horse Race Model, Verbruggen et al., 2019), reducing net activation and the resultant motor response. While the inhibitory process is rapidly engaged, its offset is less clear and was modelled to have a longer tail, approximating the offset of the preparatory process. The presence or absence of the response is determined by whether net activity crosses the initiation threshold, and its force and timing are reflected by the point at and extent to which the threshold was exceeded. With respect to the initiation threshold and how the signals are combined, a low threshold was selected, reflecting evidence from TMS and functional magnetic resonance imaging (fMRI) that inhibitory synapses are less numerous and are strategically better located than excitatory synapses, suggesting that they may be more efficient and may consume less energy than excitatory synapses (Waldvogel et al., 2000). The model shows how the variable engagement of inhibitory-control can result is a range of net activation levels that either exceed or fall below the initiation threshold by varying degrees (Fig. 6). It is important to note that while Failed Inhibitions are associated with a range of behavioural responses, the force of Successful Inhibitions is always zero and they do not have response times.

The model makes several predictions regarding changes in the level of engagement (Fig. 7). First, it predicts that P300 should be smaller on Failed compared to Successful Inhibitions. Second, it predicts that P300 should be negatively associated with force on Failed Inhibitions. Third, it also predicts a relationship between force and response time on Failed Inhibitions where larger forces should be executed earlier. Lastly, the model can also make predictions about changes in the time-course of inhibitory-control, showing that delays in the engagement of inhibitory-control of the same magnitude can result in failures to inhibit (Fig. 6B).

**Figure 7.**
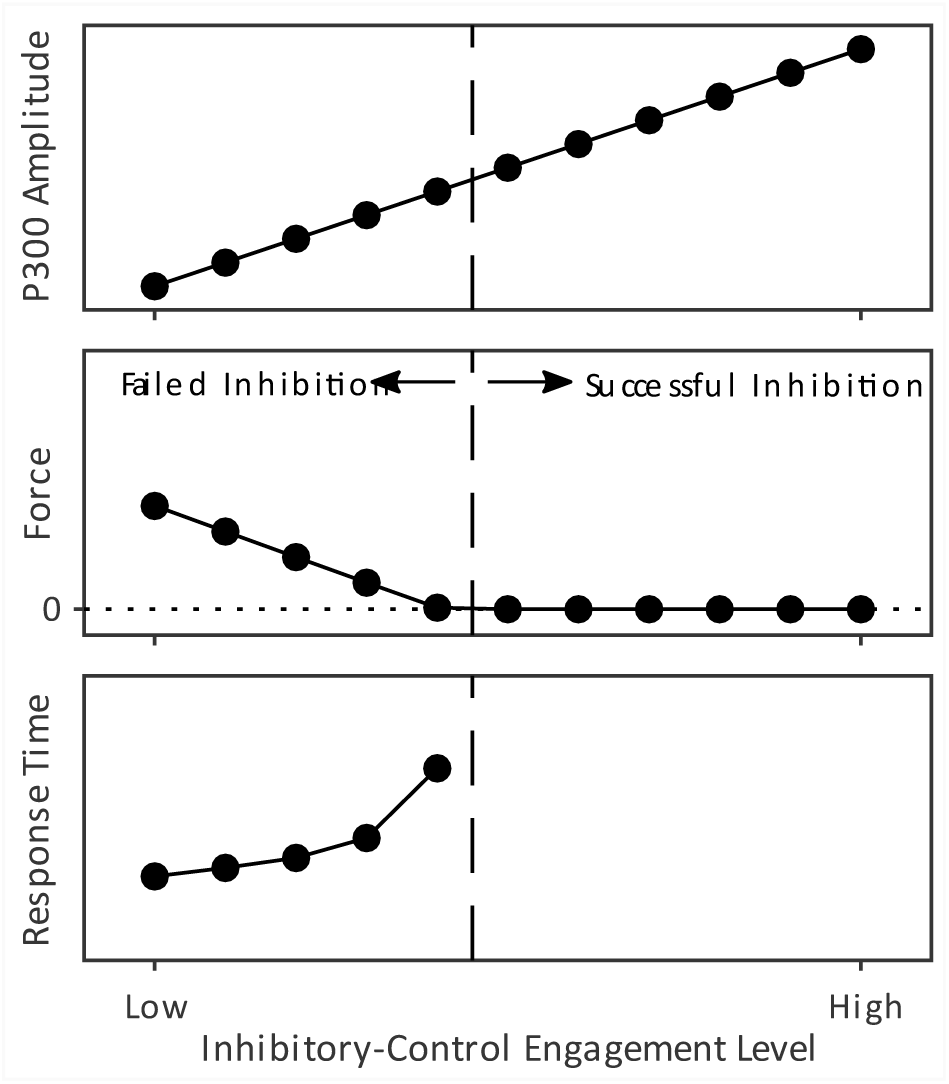
A plot depicting the expected pattern of P300 amplitude, Force and Response time as a function of inhibitory-control engagement, as predicted by the Activation-to-Threshold Model of Inhibitory-Control. As inhibitory-control engagement increases, P300 and force are shown to increase and decrease, respectively. The model shows that beyond a certain level of engagement, the response is completely suppressed and does not vary behaviourally but further increases in P300 are still expected. The model also makes predictions about the timing of responses on Failed Inhibitions.

Although we only showed examples of one-dimensional changes in the magnitude and time-course of the inhibitory-control process for illustrative purposes, it is possible for both processes to vary freely from trial-to-trial as temporal and event (e.g., the occurrence of a Stop-signal) expectancies fluctuate throughout the experiment. Additionally, we only illustrated a single inhibitory process triggered by the Stop-signal but acknowledge that additional inhibitory processes may be involved. For example, additional inhibitory processes may be engaged in non-inhibition trials to prevent the premature release of a prepared response (Duque, Lew, Mazzocchio, Olivier, & Ivry, 2010).

## 5. Conclusion

Collectively, the results provide strong evidence that the engagement of inhibitory-control varies on a continuum from trial-to-trial and that this engagement is reflected by the P300. To our knowledge, our study is the first to show continuous associations between P300 and response magnitude in the context of inhibitory-control, adding to current literature which largely focuses on P300 modulations across binary Failed and Successful Inhibition categories, and conceptualising inhibition as an all-or-none process.

Our findings have several implications for our current understanding and the future examination of inhibitory-control processing. Firstly, it highlights the utility of the Anticipatory Timing Task and the temporal control it provides over both the Stop-signal and expected response time, which allowed for the study of inhibitory-control in a high difficulty setting while maintaining a high level of control over the stimulus-environment. Secondly, it underscores the importance of studying the spectrum of motor outputs and its potential to provide further insights regarding the functional significance of other event-related brain signals (Huster, Schneider, Lavallee, Enriquez-Geppert, & Herrmann, 2017). Lastly, it draws attention to the fact that future models of inhibitory-control need to account for motor output, and not just the presence and absence of the response. Our model offers a new conceptual framework for describing the implementation of inhibition-control over highly prepared motor responses and it provides a perspective that accounts for motor output.

## Acknowledgements

We would like to thank Amy Tiberio and Emily Corti for help with data collection. The study was supported by a Discovery Project grant from the Australian Research Council (DP180100394) awarded to W.M. and O.V.L. The funding body had no role in study design, data collection and analysis, or the decision to submit the work for publication.

